# Spatiotemporal mapping of microscale stiffness during collagen polymerization and crosslinking by optical multifrequency time-harmonic elastography

**DOI:** 10.64898/2025.12.16.694603

**Authors:** Frank Sauer, Jakob Jordan, Tom Meyer, Pedro Dantas de Moraes, Guillaume Flé, Josef A. Käs, Ingolf Sack

**Author notes:** contributed equally (Frank Sauer, Jakob Jordan).

## Abstract

Optical multifrequency time-harmonic elastography (OMTHE) was used for rapid mechanical characterization of extra-cellular matrix-derived collagen networks at micrometer resolution. OMTHE was optimized for point-wise shear wave excitation in small sample volumes and compared to tabletop magnetic resonance elastography (ttMRE) and optical intensity changes. Dynamic stiffening due to the fluid-gel transition during collagen polymerization and chemical crosslinking using glutaraldehyde was tracked by shear waves speed (SWS) at vibration frequencies between 3 and 10 kHz and frame rates up to 4 kHz. During collagen polymerization, after an initial lag phase, SWS increased on average 6 ± 3 min earlier than optical density, suggesting that a load-bearing percolating fiber network was established before fibril thickening enhanced light scattering. In contrast, chemical crosslinking showed a lag-free, diffusion-driven SWS increase from 1.7 ± 0.4 m/s to 2.5 ± 0.5 m/s, matching the relative SWS change from ground-truth ttMRE. In conclusion, OMTHE provides a unique research tool that quantifies biomechanical property changes in small biological samples with spatiotemporal resolutions of micrometers and seconds.

**Key Results:**

- Point-excitation OMTHE at microscopic resolution maps dynamic stiffness changes in collagen gels during polymerization and crosslinking at high frame rates.
- Polymerization and crosslinking of collagen show distinct time courses with polymerization being in the order of minutes ahead of crosslinking.
- Collagen stiffening due to polymerization precedes changes in optical density as seen by light microscopy.

## Introduction

Biomechanical properties such as tissue stiffness are key regulators of physiological and pathological processes and provide valuable diagnostic information in vivo (1,2). The biomechanical properties of tumors are known to dynamically change as cells interact with their extracellular matrix (ECM) leading to microstructural remodeling that can be sensed by in vivo magnetic resonance elastography (MRE) (1,3). ECM-mimicking collagen gels are widely used to study basic biomechanical networks on the microscale. Various types of collagen networks are accessible for experiments in small volumes covering stiffness ranges similar to those of biological soft-tissue environments (4–6). The range of very soft properties (from a few tens of Pascals to the lower kPa-range) in combination with small sample volumes imposes experimental challenges for mechanical tests in collagen gels. To date only a few methods provide sufficient spatiotemporal resolution for tracking dynamic stiffness changes in small sample volumes at high frame rates. Table 1 lists major contemporary technologies for the quantification of biomechanical properties from the cell-scale to in vivo organs.

**Table 1:** Methodical Fragmentation in Biomechanics. Overview of current rheological measurement systems and their corresponding key parameters. Abbreviation: Optical Cell Stretcher (OCS), Real-Time Deformability Cytometry (RT-DC), Shear-Flow Deformation Cytometry (SF-DC), Atomic Force Microscopy (AFM), Optical Coherence Elastography (OCE), 7T micro MRE (µMRE), tabletop MRE (ttMRE), Oscillatory Shear Rheology (OSR), Ultrasound Elastography (USE), M_L_ longitudinal modulus, E* complex elastic modulus, k* complex wave number, G* complex shear modulus.

| Method | Frequency (Hz) | Typical probed volume/area | Typical resolution | Dynamic/Static | Measured value |
| --- | --- | --- | --- | --- | --- |
| Brillouin Mic. (20) | $10^9$ | fL | $0.3 \times 0.3 \times 1 \mu\text{m}^3$ | Static | $M_L$ |
| OCS (9) | - | pL | Cellular | Static | Deformation |
| RT-DC(10) | - | pL | cellular | Static | Deformation |
| SF-DC (11) | - | pL | Cellular | Dynamic | $G^*(\omega)$ |
| AFM (8) | 1 – 200 | $(100 \mu\text{m})^2$ | $0.1 \times 0.1 \mu\text{m}^2$ | Both | $E^*(\omega)$ |
| OCE (14) | 10 - 3000 | pL | $10 \times 10 \times 5 \mu\text{m}^3$ | Both | $c(\omega)$ |
| OMTHE (16) | 200 – 20.000 | $(0.5 \text{ mm})^2$ | $0.8 \times 0.8 \times 200 \mu\text{m}^3$ | Both | $k^*(\omega)$ |
| μMRE (21) | 200 - 6000 | 0.5 mL | $40 \times 40 \times 700 \mu\text{m}^3$ | Both | $k^*(\omega)$ |
| ttMRE (22) | 200 - 6000 | 0.5 mL | Whole sample | Both | $k^*(\omega)$ |
| OSR (23) | 0.01 – 100 | 0.5 mL | Whole sample | Dynamic | $G^*(\omega)$ |
| In vivo USE (24) | 20 – 100 | L | $(0.5 \text{ mm})^3$ | Both | $k^*(\omega)$ |
| In vivo MRE (25) | 20 – 100 | L | $(2 \text{ mm})^3$ | Both | $k^*(\omega)$ |

While MRE and ultrasound elastography methods (USE) are capable of measuring bulk stiffness properties in volume samples, tissues or organs, other methods require surface contact, transparent materials or suspended cells. However, with volume sizes in the order of mL to L, MRE and USE operate on a coarse-grained scale that does not allow microstructural assessments (7). Other methods tailored for micro-samples such as atomic force microscopy (AFM) (8), optical cell stretcher (OCS) (9), real-time deformability cytometry (RT-DC) (10), shear flow deformation cytometry (SF-DC) (11) and single cell optical elastography (12,13)required surface contact or suspended cells and are, thus, not suitable for investigating liquid-to-solid phase transitions in collagen gels. Optical coherence elastography (OCE) (14) is suitable for resolving rapid dynamic processes in transparent materials with high spatial resolution (15). However, it comes at high costs and complex laser-based experimental setups.

Therefore, we introduce optical multifrequency time-harmonic elastography (OMTHE) (16) for the investigation of polymerization and cross-linking of collagen gels. Recently, OMTHE has been introduced for static stiffness mapping in biofilms and zebrafish embryos using reverberant shear waves introduced by a variety of actuators (16). Here, we aim to extend the range of OMTHE applications to point-source wave excitation using an extended piston-driver optimized for soft gels. Piston driving allows for targeted vibrations at specific points compared to full sample excitation. In the specific case of collagen, high viscosity leads to strong damping of high frequency shear waves, making full sample excitation challenging due reduced light transmission and the capillary effect of the collagen surface within areas close to the petri dish walls. As OMTHE can map stiffness at micrometer resolutions within less than a second acquisition time, rapid polymerization and crosslinking dynamics in collagen become accessible. We previously used glutaraldehyde (GA) in collagen gels (17) to mimic fibrosis induced tissue stiffening (1,18) and measured steady-state changes by ttMRE. Due to its molecular small size, GA primarily introduces crosslinks within collagen fibers and between nearby fibrils rather than establishing wide-ranging structures (19). Therefore, GA does not alter the network structure as resolved by confocal microscopy, leaving pore size and water diffusion unaffected (18).

Aims of this study were to establish OMTHE for rapid micro-mechanical tests of liquid-solid phase transitions in soft polymer samples and quantify stiffness dynamics in collagen type-I gels during polymerization and GA-induced crosslinking in comparison to optical intensity changes and ground truth ttMRE.

## Methods

### Sample preparation and handling

#### Collagen preparation for OMHTE measurements

Collagen gels were made from a 1:2 mixture of rat tail collagen type 1 (4 g/l, Lot#230961, Serva, Heidelberg, Germany) and bovine skin collagen type 1 (4 g/l, Lot#H240502, PAN Biotech, Aidenbach, Germany) with a final collagen gel concentration of 2.5 g/l. For the preparation of 1ml gel, 0.208 ml of rat tail collagen and 0.417 of bovine skin collagen were mixed together with a phosphate buffer solution containing 0.169 ml 1M disodium hydrogen phosphate (Na_2_HPO_4_), 0.031 ml sodium dihydrogen phosphate (NaH_2_PO_4_·H_2_O) and 0.175 ml dH_2_O to achieve a final pH of 7.4 (5). To monitor collagen polymerization, the buffer and collagen solutions were mixed directly at the microscope, and 200 µl of the final mixture was immediately pipetted into a well of an µ-Slide 8 Well (ibidi GmbH, Gräfeling, Germany), which was directly transferred to incubated stage of the microscope and the measurement was started.

For the crosslinking experiments, after mixing buffer and collagen solution, 200 µl of the final solution were pipetted into each well of an µ-Slide 8 Well (ibidi GmbH, Gräfeling, Germany) and placed in an incubator at 37 °C and 95 % humidity for polymerization for at least 1.5 hours. Afterwards, 250 µl of PBS was added to each well to prevent the gel from drying out. The collagen gels were then placed on the microscope to measure them in their *native* state with OMTHE. Subsequent crosslinking was induced by removing the PBS and adding 300 µl of 0.2% glutaraldehyde solution (diluted in PBS, stock 25%, Serva, Heidelberg, Germany)(17). To ensure good sample penetration, the sample was washed with the solution two times and then incubated in it. The total time to induce the crosslinking process took less than one minute. Immediately afterwards the OMTHE time series was started.

#### Collagen preparation for MRE

Collagen preparation for ground-truth tabletop MRE (ttMRE) followed the same steps as described above but with larger sample volumes of 500 µl in order to provide sufficient material for 7-mm diameter glass tubes used in the ttMRE experiments. After incubation of collagen samples for 1.5 hours at 37 °C, gels were investigated in their *native* state. For crosslinking, 1 ml of the 0.2% glutaraldehyde solution described above was added for one hour. Afterwards, the gels were washed 5 times with PBS and subsequently investigated in their *crosslinked* state.

### Tabletop MRE

A 0.5-Tesla permanent-magnet MRI scanner (Pure Devices GmbH, Würzburg, Germany) with 10-mm bore, external gradient amplifier and a piezoelectric driver was used (18,22). The sample was placed in a glass tube of 7 mm diameter and kept at 37 °C. Concentrical shear waves were excited via the glass wall of the sample tubes primarily along the cylinder axis. 5 samples of collagen type-1 before and after crosslinking with glutaraldehyde were investigated at frequencies *f* = 250, 400, 520, 700, 850, 1000 and 1150 Hz. For each frequency 8 raw-phase images were unwrapped and Fourier-transformed in time to extract the complex-valued fundamental frequency. Radial wave profiles were fitted by an analytical solution of shear waves in a z-infinite cylinder (22), providing the complex wave number *k* = k’ + ik’’*. Shear wave speed *c* and the shear wave penetration rate *a* were derived from *k\** for each frequency according to

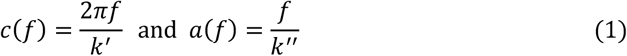

The complex shear modulus *G* =* |*G\**|·*e*^*iφ*^ can then be derived from *c* via

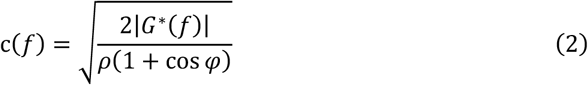

with unit density *ρ* = 1 kg/l, which roughly matches most soft tissues (26). Viscoelastic dispersion was fitted with a generalized fractional Maxwell model (27), which has shown good applicability on both eukaryotic cells (28) and biological tissues (27),

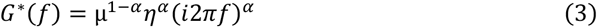

This model can interpolate between a pure elastic solid with the spring constant *µ* resembling *G’* for a power law variable *α* → 0 and a viscous fluid with the dashpot parameter *η* for *α* → 1 (27). Since *µ* and *η* are linearly dependent, this 3-parameter model was reduced to a 2-parameter model by setting *η* to 1 Pa·s. By multiplying *α* with π/2, it can be directly translated into the phase angle *φ* of the complex shear modulus *G\**. As the parameters from this viscoelastic fit model are more robust than data obtained from single frequency values, we used these for comparison with OMTHE, by calculating a comparative shear wave speed c at 6333 Hz, which represents the center of the OMTHE frequency range. Consequently, Eq. 2 and 3 can be combined to express the fractional KV model in *c* and *a* notation:

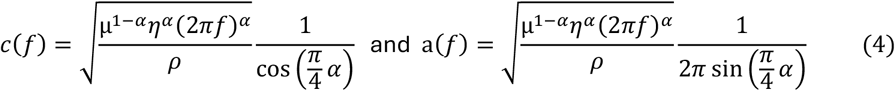

### OMTHE measurements

Figure 1. shows the setup of OMTHE for collagen samples investigations. Vibrations were induced using a needle-tipped piezo actuator (APA35XS, Cedrat Technologies, France), driven by a piezo actuator amplifier (PX200, PiezoDrive, Australia) with the frequency being controlled by a frequency generator (AFG 3022B, Tektronix, USA). The same frequency generator also triggered a high-speed camera (Fastcam Mini AX100 type 540K-S, Photron, Japan) to record optical intensity images with 1024 x 1024 pixels at an ISO of 40 000. Image acquisition was synchronized with piezo vibration (Supplemental Video S1), to stroboscopically capture shear waves. The camera was mounted on an inverted microscope (Axio Observer, Zeiss, Jena, Germany) equipped with an incubation stage (37 °C and 95 % relative humidity) and a 25x water immersion objective (LCI Plan-Neofluar 25x/0,8 Imm Korr DIC M27, Zeiss, Jena, Germany) yielding a pixel size of 0.8 by 0.8 µm. Focal plane was 150 µm deep in the gel with the actuator needle placed within the focal plane using a hydraulic micromanipulator (Leitz, Wetzlar, Germany). Vibrations were consecutively induced at 3, 4, 5, 7, 9 and 10 kHz, omitting 6 kHz and 8 kHz due to system resonances. For each frequency, 64 images were recorded supporting 8 wave dynamics over 8 vibration periods. Total measurement time for all 6 frequencies was less than a second while transferring the total of 395 images to the workstation took approximately 30 seconds. For the collagen polymerization experiments, timepoints were set at 5 min intervals, while during the lag phase and following increase in SWS and OD it was reduced to 1 min. In contrast, collagen crosslinking was tracked at fixed timepoints of 0, 1, 5, 10, 15, 30, 45 and 60 minutes, where the first time point was assigned to the *native* state immediately followed by the addition of GA.

**Figure 1:**
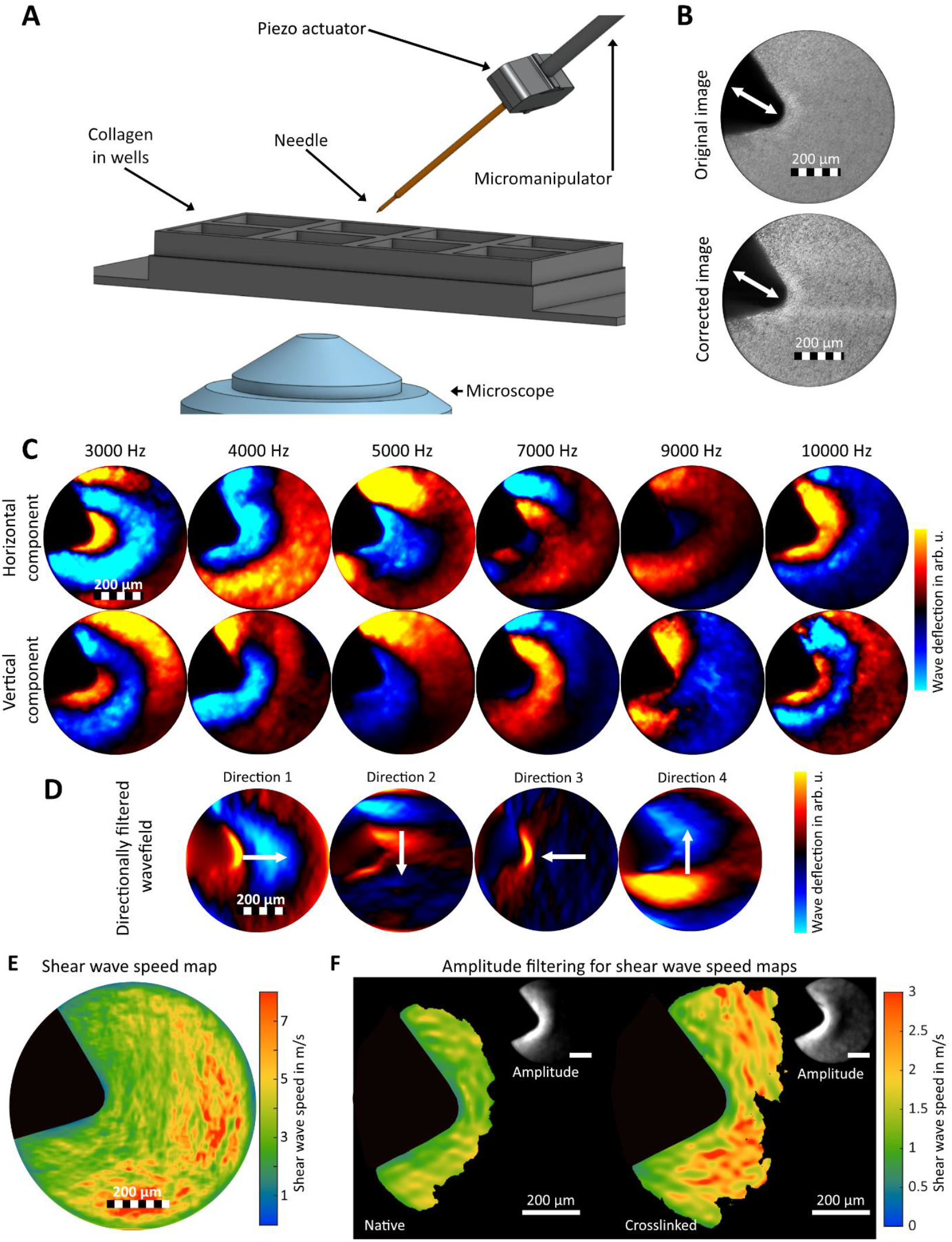
OMTHE in collagen samples using localized excitation. **(A)** Point-wise shear wave excitation using a 1 mm metal tip on an inverted microscope. **(B)** Vibration-synchronized brightfield images of the rod tip in contact with collagen gel before (left) and after (right) image pre-processing. White arrows indicate vibration direction. **(C)** Corresponding wave fields detected via optical flow between 5000 and 10000 Hz. **(D)** Directionally filtered wavefield during k-MDEV inversion. **(E)** SWS map of collagen gel reconstructed using k-MDEV pipeline. Viscoelasticity leads to SWS dampening and results in abnormally high SWS values in areas with minimal wave amplitude further away from the tip. **(F)** Amplitude filtering to counter the behavior described in **(E)**. Based on the magnitude of the complex wave amplitude (see insets) we defined a cutoff threshold of e^-1^. The resulting SWS maps including manually drawn tip masks are shown for an example of Native and GA Crosslinked collagen gel. Due to the transition towards a more solid-like or elastic material the crosslinked collagen gels exhibit a decreased shear wave dampening (compare also ttMRE wave fields in **Figure 2 Figure 2** A and B and as consequence also a larger ROI after amplitude filtering.

### Data evaluation

#### Optical Density

Optical Density (OD) was measured as brightness decrease based on raw images in .tif-Format before further preprocessing. For each sample and its corresponding measurement series a manually drawn ROI was used to mask out the borders of the well plate and the metal tip from the FOV. Consequently, the intensity data from the remaining FOV was averaged, OD calculated as 1 – image brightness and normalized to the maximum value at the end of the timeseries using a custom built matlab code (Matlab version R2024a, The MathWorks Inc USA).

#### Motion estimation

To correct light flickering the image series for each frequency was normalized to the median of their central zone. Motion correction was then applied to remove jitter and reduce unwanted rigid motion followed by histogram equalization to linearize the dynamic intensity range (16). A harmonic optical flow algorithm was applied to estimate xy-period displacements (**Figure 1Figure 1**C and Supplemental Video S2) at fundamental frequency as described in (16).

#### Parameter reconstruction

In previous work, OMTHE was based on direct multifrequency inversion (MDEV) (16). In this study, we expanded the multi-inversion pipeline to noise-robust wavenumber inversion (k-MDEV) which is used in MRE and ultrasound-based time harmonic elastography (29–31). Retrieval of wave numbers by k-MDEV requires spatiotemporal filters to avoid phase gradient disturbances by wave interferences. Spatiotemporal filters were applied along the four principal directions of the xy-coordinate system in k-space (**Figure 1**D)**Figure 1**. According to (32), linear high-pass filters with a Butterworth-low pass of thresholds of 0.001 and 0.05 m^-1^ were applied for each frequency, wave component and wave direction prior to the derivative of phase-gradients. All phase gradient images were averaged and inverted to compound OMTHE SWS maps (**Figure 1Figure 1**E). The parameters were evaluated within regions of high amplitudes (> e^-1^ with reference to xy-amplitudes at wave source near the driver tip) to minimize effects of critical attenuation (**Figure 1Figure 1**F).

#### Polymerization kinetics

To compare the kinetics of SWS and OD, relative changes were analyzed. For this purpose, each SWS measurement series was normalized to the mean of the final three time points, while the OD data were normalized to the last time point. Mean and standard deviation were calculated for both normalized and SWS and normalized OD. To model the characteristic dynamic of collagen polymerization (33) consisting of a initial lag phase that is followed by an increase to a plateau value we used a time-shifted Avrami model (34):

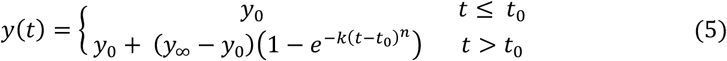

where y_0_ is defined as the average of the first 3 time-points as baseline during the lag phase before quantifiable parameter change. When the increase surpasses 2· SDs of *y*_*0*_ the onset time *t*_*0*_ is reached. *y*_∞_ represents the plateau values. *k* is the rate constant and *n* as the Avrami exponent reflects the dimensionality and cooperativity of the polymerization process with *n* ≈ 1 for one-dimension or purely diffusion-limited growth, while higher values indicate multidimensional network formation(35). By further scaling with

Eq.5 can be reduced to:

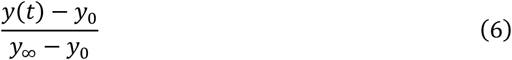

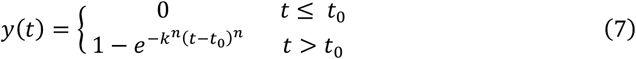

The characteristic half time *t*_*50*_ can be derived as:

#### Crosslinking kinetics

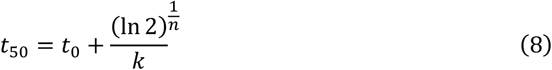

In contrast to the polymerization data, the temporal evolution of SWS during additional crosslinking followed first-order kinetics and could be fitted with a single-exponential approach to a plateau:

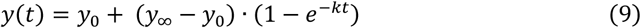

Individual time series were normalized to the first value assigned as t = 0 min before the addition of GA, so that *y*_*0*′_= 1 was the *native* reference. t = 1 min after GA was taken as the first time point for the fit. Plateau value *y*_∞_ was calculated as mean of the last three time-points. The characteristic half time for the crosslinking experiments t_*50*_ followed

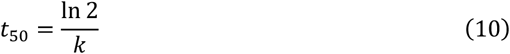

## Results

### Tabletop MRE as ground truth measurements

Typical measurement times for a multifrequency ttMRE experiment were in the range of one hour, rendering dynamic measurements on the time scale of collagen polymerization and crosslinking impossible. Therefore, we used ttMRE as a ground truth modality to mechanically quantify *native* and end-point crosslinked states. Representative wave fields for all frequencies in both conditions are shown in **Figure 2**A. Wavelengths decreased over frequency while GA crosslinking increased wavelengths and reduced wave attenuation as demonstrated by the radial wave profiles in **Figure 2**B for 400 Hz and 1150 Hz. Viscoelastic dispersion is shown in **Figure 2**C where dashed lines represent springpot-fits according to Eqs.2 and 4. During crosslinking the power-law parameter *α* decreased from 0.45 ± 0.06 to 0.31 ± 0.02, while shear modulus *μ* increased from 49 ± 34 Pa to 587 ± 50 Pa. Using Eqs. 2 and 3, SWS was derived for a hypothetical frequency of 6333 Hz - the center of the OMTHE frequency range - showing an increase from 1.03 ± 0.05 m/s to 1.52 ± 0.07 m/s, which represents a relative increase of 47 ± 9% (**Figure 4**C).

**Figure 2:**
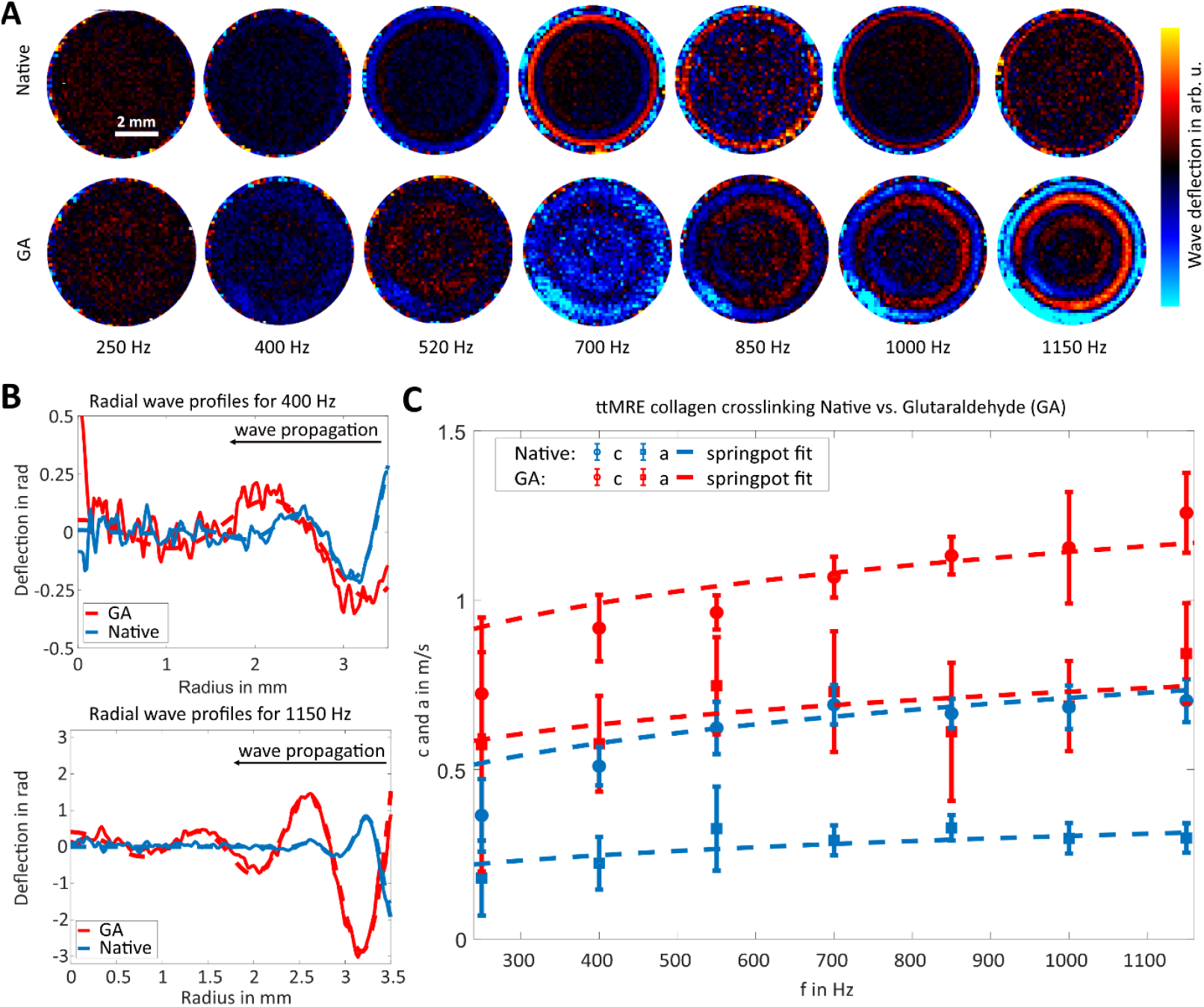
Ground truth measurements with ttMRE: Collagen crosslinking with glutaraldehyde. **(A)** Representative ttMRE wave fields for native (top row) and GA crosslinked collagen (bottom row) over the examined frequency range from 250 Hz to 1150 Hz. Same scaling across all images. **(B)** Radial wave profiles for 400 Hz (top) and 1150 Hz (bottom). Data extracted from the wave fields depicted in (A). For better visualization only the real part of the wave signal is shown (solid thin lines). To extract k*, the wave propagation was analytically fitted (thick dashed lines). Wavelength increased with frequency, while GA crosslinking led to longer and less dampened shear waves. **(C)** Shear wave speed c and shear wave penetration rate a increased after GA crosslinking (mean ± SD, N = 5). Dashed lines show springpot model fits according to Eq. 4.

### OMTHE during collagen polymerization

Polymerization transformed liquid collagen solutions into solid gels. This mechanical phase transition, captured by OMTHE shear waves, was accompanied by a transition of the optical properties from a clear gel to the opaque solid, captured by OD measurements. **Figure 3**A shows brightfield optical images and OMTHE shear wave amplitudes for selected timepoints during collagen polymerization. The full timeseries is shown in Supplemental Video S3. Image brightness was normalized to the last timepoint and plotted over time for 8 samples in **Figure 3**B. The curve shows an initial lag phase, which is then followed by a step increase towards a plateau. The corresponding SWS evaluations are shown in **Figure 3**C. The observed SWS variance among the five different samples was reduced by normalizing individual SWS curves to their plateau values reached at the last three time points (group mean plateau SWS: 1.6 ± 0.5 m/s). **Figure 3**D shows the normalized and group-averaged SWS and OD curves indicating a spread in the onset time of gel polymerization.

**Figure 3:**
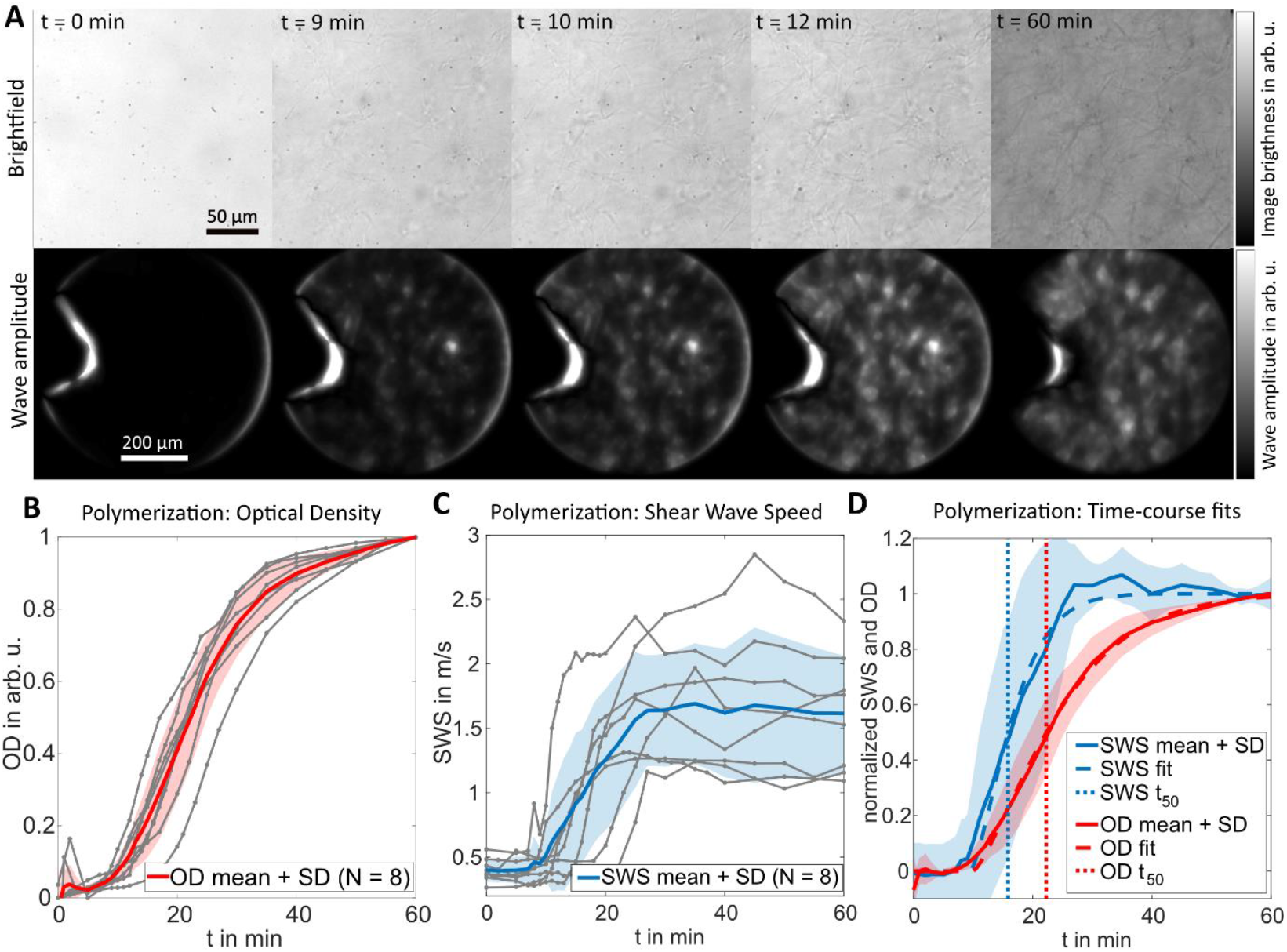
Collagen polymerization. **(A)** Time-lapse series of zoomed-in OMTHE brightfield images (top) shows the onset of collagen network formation after 9 min. Corresponding wave amplitude maps (bottom) depict signal increase over the same time frame. **(B)** Normalized optical density (OD) increased during collagen polymerization. Analysis based on intensity values from brightfield images. The onset of the OD increase varies in between samples. **(C)** Dynamic OMTHE measurements during collagen gel polymerization with the setup from Fig. 1. After an initial lag phase the SWS increased towards a plateau at (1.6 ± 0.5) m/s. **(D)** Normalized SWS and OD from 0 to 1 and corresponding Avrami fits show similar onset t_0_but distinct half times t_50_ for SWS and OD, with OD t_50_ being 6 ± 3 min slower than SWS t_50_.

To quantify polymerization kinetics, mean normalized data was fitted with the Avrami model (Eq. 5) resulting in good fits for SWS (*k* = 0.13 ± 0.07, *n* = 1.4 ± 0.9, R^2^ = 0.99) and OD (*k* = 0.06 ± 0.02, *n* = 1.3 ± 0.6, R^2^ = 0.99). While no difference was observed between the lag times t_0_ of individual samples for SWS (11 ± 4 min) and OD (10 ± 3 min, p = 0.79), the corresponding half-times t_50_ differed between SWS (17 ± 6 min) and OD (22 ± 3 min, p = 0.042), suggesting that the processes underlying SWS changes proceed on average Δt_50_ = 6 ± 3 min faster than those underlying OD changes, even though both start at the same lag time.

### OMTHE during collagen crosslinking with glutaraldehyde

OD analysis showed that collagen crosslinking had no detectable effect on optical intensity images (**Figure 4**A). In contrast, OMTHE based SWS increased towards a plateau. Similar to the effect of polymerization on SWS (**Figure 3**C), a wide variance in absolute SWS was observed among the 5 investigated samples. Average SWS increased from 1.7 ± 0.4 m/s to a plateau value of 2.5 ± 0.5 m/s. Normalized to the first (native) timepoint a 1.5 ± 0.3-fold increase was observed (**Figure 4**C) in agreement to the 1.5 ± 0.1-fold increase observed with ttMRE (**Figure 3**C). Absolute SWS values were not correlated with relative SWS changes (r = 0.27, p = 0.67).

According to Eq. 9, the time course analysis of normalized SWS followed a single exponential increase, with a rate constant *k* = 0.089 ± 0.013 min^-1^ and a characteristic half time *t*_*50*_ = 7.9 ± 1.1 min.

**Figure 4:**
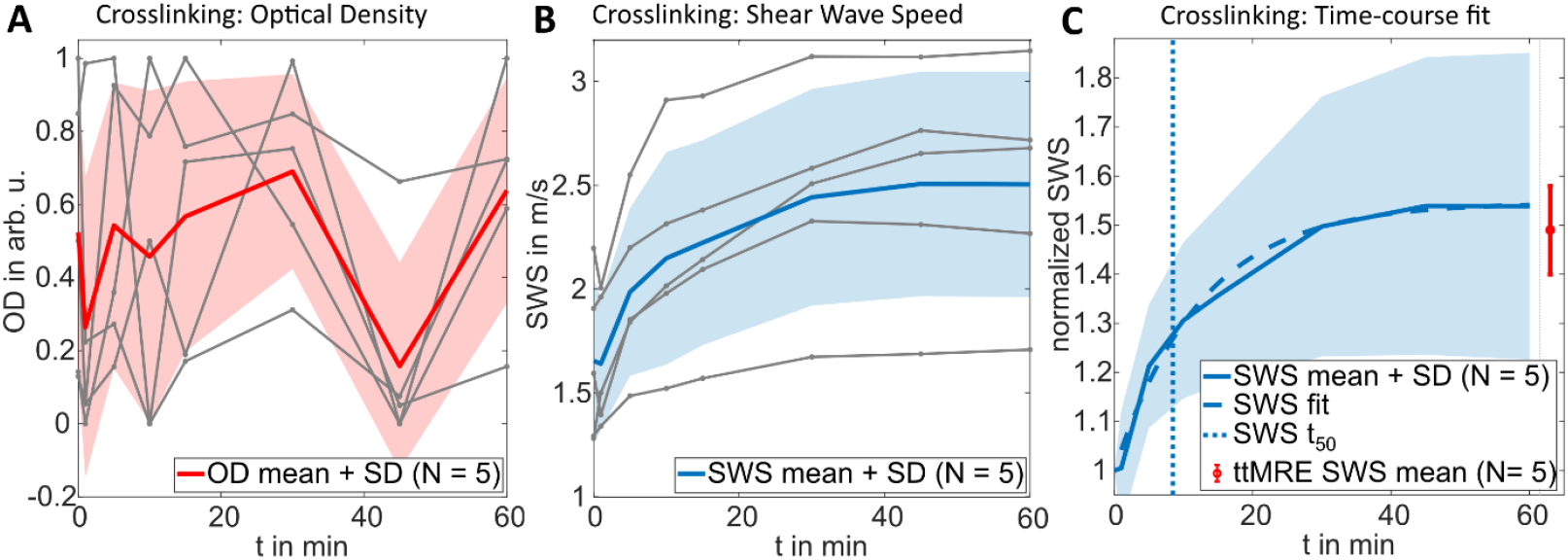
Collagen crosslinking with glutaraldehyde. (A) Normalized OD analysis during collagen crosslinking with 0.2 % glutaraldehyde. No trend is observed. (B) Dynamic OMTHE measurements showed an increase in SWS following application of crosslinker at t = 1 min. On average (N = 5), SWS rose from 1.7 ± 0.4 m/s to 2.5 ± 0.5 m/s. (C) Normalized SWS changes showed on average 1.5 ± 0.3-fold increase (N = 5) with a half rise time t_50_ of (8 ± 1) min indicating a predominantly diffusion-driven crosslinking process. Tabletop MRE measurements (red inset) under identical gel conditions showed a comparable 1.5 ± 0.1-fold increase (N = 5).

Assuming that GA diffused into the gel sample from the top surface, the diffusion time of GA could be estimated with *t*_*diff*_ =4 *L*^2^ /(ϖ^2^*D*). 200 µl of collagen gel in a rectangular well of 9.4 x 10.7 mm^2^ size resulted in *L*= 2 mm height. We have shown previously that the diffusion coefficient *D* is very similar in collagen gel and free water (18,36) with approximately 3.0·10^-9^ m^2^s^-1^ at 37 °C (37) . The resulting GA diffusion time *t*_*diff*_ = 9.0 min, is in good agreement with *t*_*50*_ reported above, indicating that the crosslinking process of collagen by GA is controlled by the diffusive transport of the cross-linker to the collagen chains.

## Discussion

By dynamically probing mechanical alterations in collagen hydrogels during polymerization and crosslinking, we demonstrate the sensitivity of OMTHE for time resolved mapping of mechanical changes in microscopic systems, while employing the same wave excitation and inversion processes as MRE und USE to enable direct comparison, interpolation and interpretation of SWS across scales.

The collagen gels used here were viscoelastic materials with a significant viscous contribution to their complex shear modulus. This becomes clear by looking at the powerlaw parameter *α* from the springpot fits in the ttMRE measurements described above. Translated into *storage* (*G’*) and *loss modulus* (*G’’*) as equivalent for *elastic* and *viscous* material properties, *native* collagen gels have a *G’’* of 0.85·*G’*, while in the *crosslinked* state it is still 0.53·*G’*. Our indenter OMTHE (**Figure 1**) acts as a point-like source for shear wave excitation where the radially emanating waves are attenuated by geometric loss. In addition, viscous damping exponentially damps the wave amplitudes with distance from the wave source. To account for these two superimposed wave attenuation processes and to avoid insufficient wave penetration, we applied an empirical amplitude threshold of e^-1^ beyond which we disregarded SWS values. The obtained SWS values during collagen crosslinking were consistent with ground truth ttMRE values and previous work (18). Notably, polymerization kinetics measured by SWS and OD showed initial lag phases followed by steep increases towards plateau values with similar onset times *t*^*0*^ of 10 ± 4 versus 10 ± 3 min.

Also the Avrami fits of mean data yielded similar exponents (OD: n = 1.3 ± 0.6, SWS: n = ± 0.9), indicating slow, weakly time-dependent formation of initial fibril nuclei followed by mainly one-dimensional fibril extension, as expected from Avrami theory for exponents near 1–2 (38). The higher rate constant and shorter half-time in the SWS measurements (k = 0.13 ± 0.07; t_50_ = 17 ± 6 min) compared to OD (k = 0.06 ± 0.02; t_50_ = 22 ± 3 min) indicate that mechanical changes within the forming network is established earlier than the OD increase. This agrees with the observation that a continuous fibrillar network is already formed while turbidity, i.e. light scattering, remains low while main OD increases occurs later during fibril thickening (39). Forgacs et al. (40) have shown sigmoidal G′ curves with an initial lag phase with oscillatory shear rheology, while separate imaging revealed early branching structures (∼60 s) that grew into larger clusters intersecting after ∼10 min, at which point G′ rose sharply. Bulk OD measurements using the same preparation reported a ∼30 min lag time [23]. Yang et al. (41) further identified an arrest time, marking cessation of cluster motion, coincided with or slightly preceded by the onset of G′ increase. Together, these studies show that fibrils appear early, while mechanical rigidity emerges only when a percolated network forms, consistent to the early rise of SWS ahead of OD in our study. OD reflects late fibril thickening by light scattering that is integrated over the entire sample thickness. Together, our findings support a unified picture of collagen assembly: early appearance of thin fibrils, gradually slowing nucleation, continued elongation of existing fibrils, and earlier mechanical percolation relative to the later rise in turbidity.

The sensitivity of OMTHE to rapid mechanical changes is also evident during GA crosslinking, where OD, as expected, showed no detectable change. GA acts at the nanometer scale as a strong inter- and intrafibrillar crosslinker and does not generate new structures or alter pore size (18), making optical measurements largely insensitive to its effects. Both the bulk ttMRE and microscopic OMTHE samples were exposed to an excess of GA solution, resulting in comparable relative increases in stiffness (1.5 ± 0.1-fold vs. ± 0.3-fold, respectively). Although time-resolved mechanical data during GA treatment are limited, existing studies consistently report an immediate stiffening response. Ho et al. (42) observed an instantaneous rise in viscosity followed by a plateau after ∼20 min at the GA concentration used here, and Nam et al. (43) similarly reported a rapid increase in elastic modulus toward a plateau. Comparable kinetics have been described for genipin crosslinking (44). These reports agree with our observations, conforming that OMTHE detects the fast diffusion-driven mechanical stiffening upon GA addition, that optical readouts miss.

## Limitations

Although the nominal frequency ranges of ttMRE (200–6000 Hz) and OMTHE (300–10000 Hz) overlap, practical constraints in wave propagation, geometry, and FOV create an effective gap between 1200 Hz and 3000 Hz, preventing direct comparison. After GA addition, OMTHE SWS immediately increased towards plateau values with relative changes similar to those obtained by ground-truth ttMRE. However, springpot-based extrapolation (Eq.3) of ttMRE data still underestimated absolute SWS, indicating that one parameter set cannot sufficiently predict the SWS dispersions of both frequency ranges of OMTHE and ttMRE. OMTHE curves also showed spread in absolute SWS values and, during polymerization, in onset times (**Figure 3**C and **Figure 4**B). Batch-to-batch differences in collagen handling, individual sample preparation, and the small OMTHE focal volume likely contributed, as did manual indenter positioning and sensitivity to local inhomogeneities. The oscillating tip with amplitudes in the order of 10 µm may have perturbed network formation by providing additional nucleation sites, leading to earlier onset times than bulk turbidity (33). However, this potential perturbation would also apply to other dynamic probing techniques, including AFM and OSR. Early SWS values were further limited by the initially small FOV imposed by amplitude filtering. In the GA crosslinking measurement series the network was already formed and then progressively stiffened (1,18), improving wave propagation and data quality.

## Conclusion

In summary, this study demonstrated that OMTHE optimized for small polymer samples using a point-wave source enables rapid, time-resolved characterization of reaction kinetics in ECM-mimicking collagen gels. The method captured key mechanical transitions from liquid to solid phases during both polymerization and crosslinking. Distinct kinetic phases and characteristic times were observed for polymerization based on stiffness changes and optical intensity changes, suggesting that mechanical percolations of fibrillar networks preceded optical thickening while the observed stiffening through crosslinking was diffusion-driven. In this context, OMTHE may also be a useful alternative to AFM and related techniques for in vitro systems that focus on dynamic mechanical changes, such as disease-induced matrix remodeling (45). Together, our findings offer insights into polymer dynamics of soft biological gels and pose OMTHE as a useful research tool for localized mechanical measurements during rapid biochemical processes at micrometer resolutions.

## Video appendix

S1 tip collagen brightfield S2 animated wave fields

S3 amplitude change polymerization

