## Supplementary figures and images for "Spatiotemporal mapping of microscale stiffness during collagen polymerization and crosslinking by optical multifrequency time-harmonic elastography"

### Supplemental Video S1

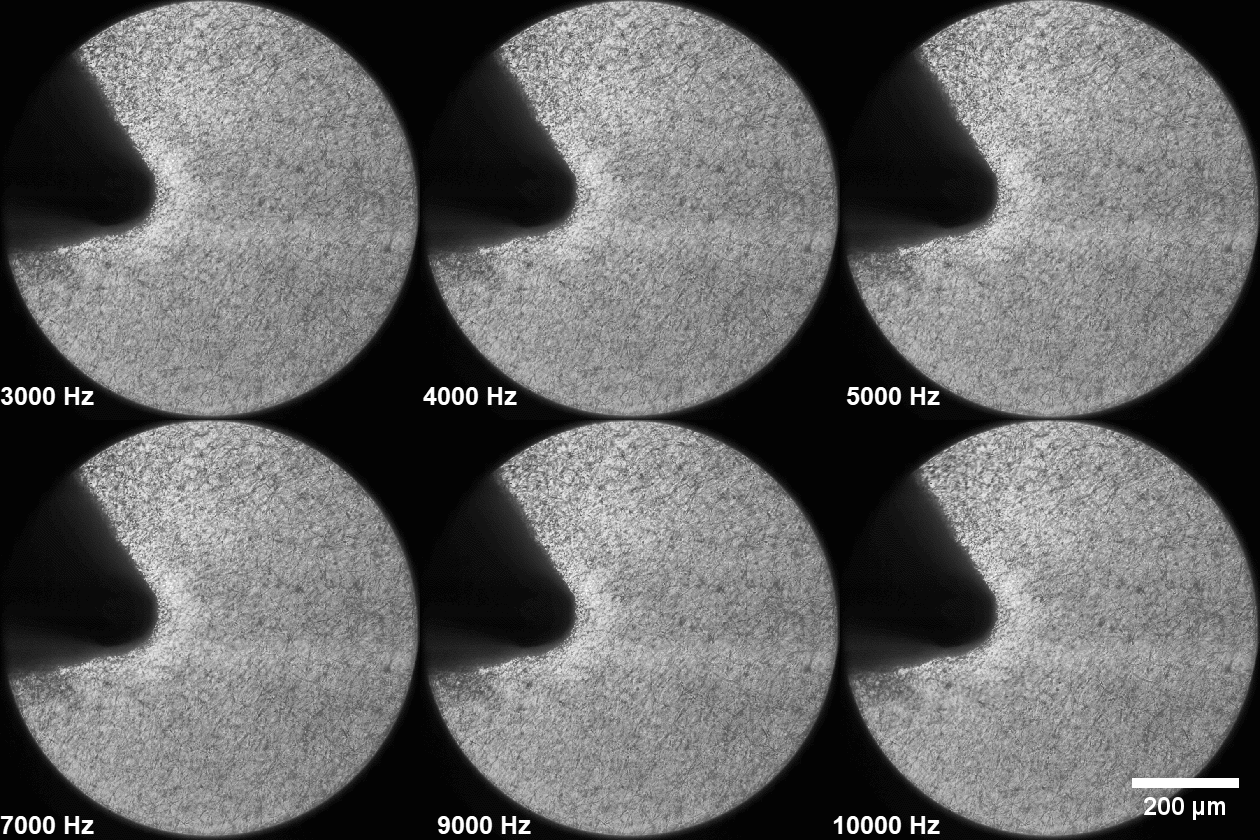

### Supplemental Video S2a

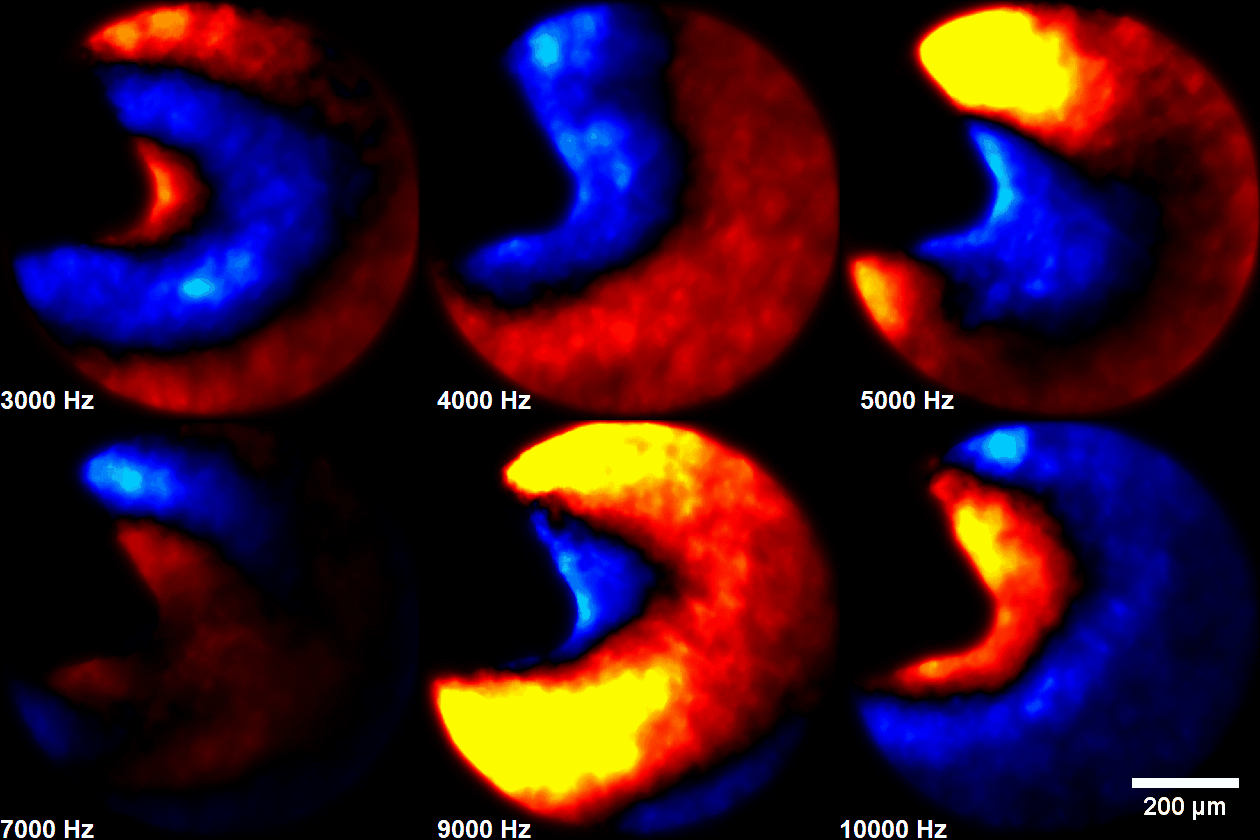

### Supplemental Video S2b

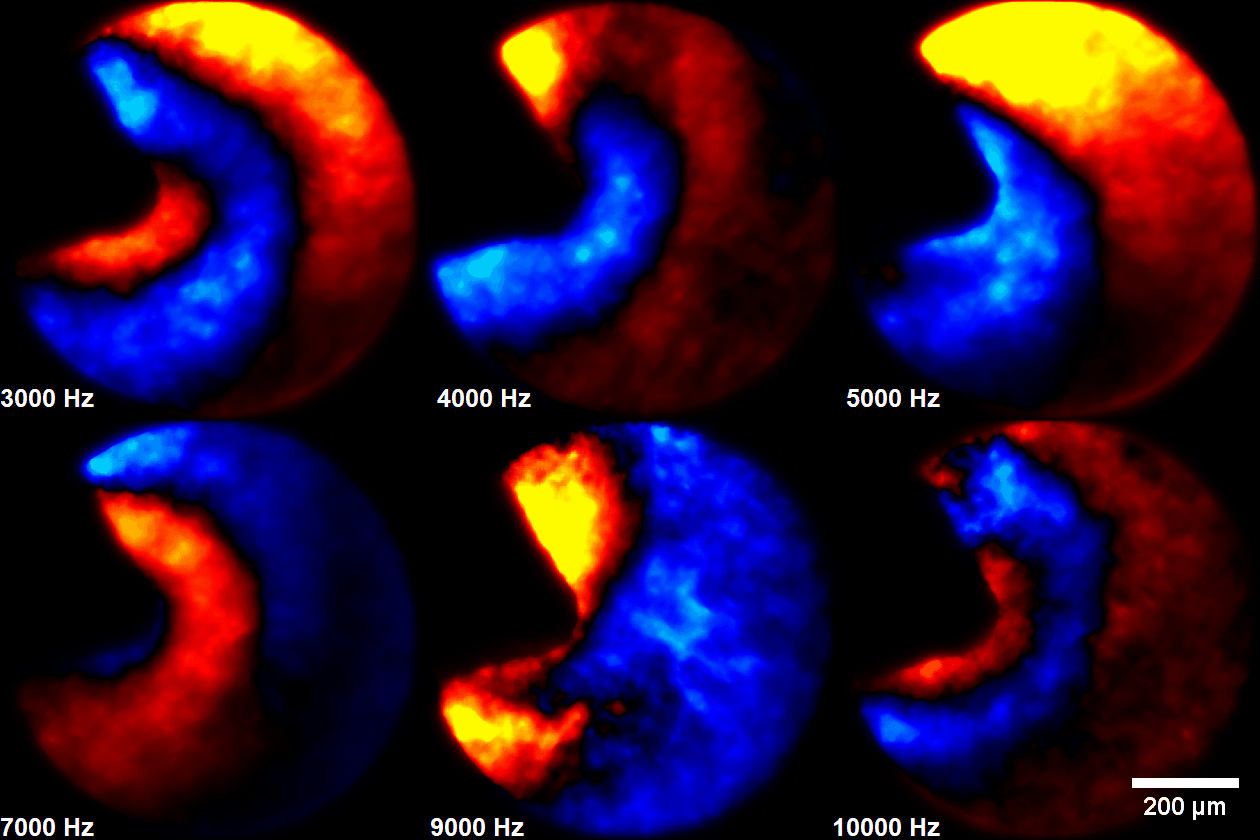

### Supplemental Video S3

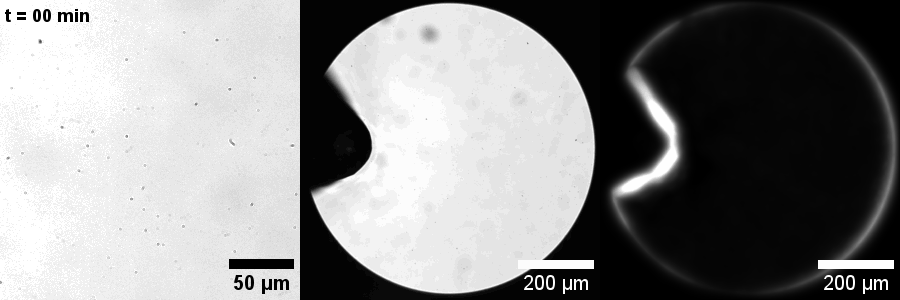
